# DEVIL peptides control cell growth and differentiation in different developmental processes

**DOI:** 10.1101/2023.09.13.557675

**Authors:** Ana Alarcia, Amparo Primo-Capella, Elena Perpiñán, Priscilla Rossetto, Cristina Ferrándiz

## Abstract

The DEVIL/ROTUNDIFOLIA-LIKE (DVL/RTFL) family of plant peptides is present in all land plants, but their biological role and mode of action remains largely unknown, in part due to the lack of reported phenotypes associated to DVL/RTFL loss of function. In this work we have generated high order mutants and characterized their phenotypes in reproductive development. Our results indicate that *dvl* mutants are affected in cell elongation processes, mainly related to pollen tube growth, and they appear to provide robustness to these processes. We also show that DVL peptides may act in different domains to those where the corresponding genes are transcribed, suggesting a putative role in the coordination of organ growth by participating in cell-to-cell communication.

## Introduction

Peptides are usually recognized as proteins smaller than 100 aminoacids, a generally accepted limit, however arbitrary[1]. Plant peptides play a wide range of functions in plant morphogenesis, fertilization, cell-to-cell signaling, triggering cell death and senescence, and integrating environmental signals, and are key players in defense responses to biotic stress or in interorganismal communication [1–4]. This functional diversity is also reflected in the huge differences found in their structural features and mode of action, that can be mediated by secretory properties that allow messenger roles, by the interaction with specific receptors that elicit downstream signaling cascades or by direct interference with the transcription and translation machinery to reprogram overall cell functioning[1–5]. Plant peptides can be classified into different categories depending on their origin, if they are derived from processing of nonfunctional precursor proteins or functional proteins (acquiring then a different function from the precursor), or if they are directly translated by small open reading frames (sORFs) [1,2]. While most of the functionally characterized plant peptides are derived from precursors, the number of known peptides derived from non-precursors is growing, although only a handful has been functionally characterized. Among these peptides encoded by sORFs, the DEVIL/ROTUNDIFOLIA-LIKE (DVL/RTFL) family, which in Arabidopsis is comprised by 24 members, has been known for many years, although their precise biological functions remain a mystery, in part because loss-of-function mutants characterized so far have not shown any associated phenotypes [6,7]. The DVL/RTFL family is present in all land plants, to which is specific [8,9]. In sequenced genomes of species in most plant taxa the number of members of the family that can be identified is generally low. For example, only a single DVL gene has been found in *Marchantia polymorpha*, two paralogs in *Physcomitrella patents*, 2 to 5 in sequenced gymnosperm species and two paralogs in the basal angiosperm *Amborella thricopoda* [9]. However, the family has expanded significantly through angiosperm evolution and around 20 family members can be found in available high-quality genomes of both monocot (such as *Oryza sativa*) and dicot species (*Arabidopsis, Solanum lycopersicum, Pisum sativum*…), suggesting that functional diversification must have occurred in flowering plants [9].

DVL/RTFL genes were initially identified in Arabidopsis by the phenotypes observed in two independent activation-tagged lines: *devil1-1D* (*dvl1-1D*;[7]) and *rotundifolia4-D* (*rot4-1D*; [6]. Both mutants showed similar developmental phenotypes, with round and short rosette leaves, stalked trichomes, compact inflorescences, short floral organs, and alterations in fruit shape. Overexpression of any other member of the family produce similar phenotypes in leaves, flowers, and overall plant architecture, although some differences are observed in the final shape of the siliques, indicating that they share a common mode of action [6,7,10].

DVL/RTFL peptides have a conserved region of 32 aminoacids, the RTF domain, that has been shown to be sufficient and essential to produce these constitutive expression phenotypes [11]. Most of the DVL/RTFL peptides also have highly variable N-terminal or, less frequently, C-terminal extensions that might be related to their specific functions, but that have not been characterized in detail [9]. DVL/RTFL peptides are localized in the plasma membrane, and the experimental evidence gathered so far indicate the DVL/RTFL peptides are not secreted or post-translationally processed, appearing to act locally[11]. The characterization of the overexpression lines in Arabidopsis suggests that DVL/RTFL peptides could provide positional cues to change cell proliferation patterns that could be linked to organ shape control [6]. In *Medicago truncatula*, a DVL/RTFL genes, *MtDVL1*, has been associated with nodule formation, and *M. truncatula* roots overexpressing *MtDVL1* show a reduced number of nodules, decreasing the cell cycle reactivation produced by rhizobia infection, and again pointing to a role of DVL/RTFL peptides in cell proliferation [12].

The similarity of the overexpression phenotypes of the members of the family, even in heterologous systems (like for example in Arabidopsis lines overexpressing rice paralogs [9]), and the lack of phenotypic defects caused by loss-of-function of DVL/RTFL genes [6,7] suggest that there is a high degree of functional redundancy within the family. On the other hand, this likely redundancy has hindered the elucidation of DVL/RTFL roles and mode of action. In this work, we have embarked in the generation and characterization of *dvl* mutants to provide new evidence that could help to understand the bases of their biological function.

## Results

### High order combination of *dvl* mutations cause alterations in transcriptome expression

DVL/RTFL peptides have been proposed to have a role in the coordination of growth polarity based on reported phenotypes conferred by the overexpression of some of the members of the family [6,12]. However, the lack of loss-of-function phenotypes has made difficult to characterize their biological function. To clarify the role of these peptides in development, we have performed diverse experiments that provide new evidence towards this goal.

We generated multiple combinations of mutants by genetic crosses together with genome editing mediated by CRISPR-Cas9 [13]. First, and considering the likely functional redundancy that could be encountered, we targeted members of a defined clade of the family composed of five genes and we made use of T-DNA insertion mutants available in public collections for *DVL8* (Salk_09633C), *DVL11* (Salk_204467), *RTFL9* (Salk_122220.20), *RTFL11* (GK_321A11) and *DVL19* (Sail_691_G09). The position of the T-DNAs was confirmed by PCR and sequencing of the corresponding genomic fragment for each mutant (S1 Fig A). Except for *DVL8*, where the T-DNA insertion was interrupting the conserved DVL/RTF functional domain, the insertions were positioned downstream these sequences or in the 3’UTR. Therefore, to assess the potential impact of the T-DNA insertion in the function of the gene, we quantified the expression of the corresponding genes by qRT-PCR in the mutants. In most cases, the insertion caused the reduction of the transcription of the genes by 80% or more, except in the case of *RTFL9*, where transcription was increased (S1 Fig B). We then considered *bona fide* loss-of-function alleles all but Salk_122220.20, affected in *RTFL9*, and generated double, triple, and quadruple mutant combinations, none of which showed any conspicuous defects in development. We then used CRISPR-Cas9 genome editing on the quadruple mutant to target independently *RTFL9*, *DVL1* and *DVL4*, aiming to cause deletions encompassing a significant portion of the CDS of the corresponding genes. We selected homozygous plants where the deletion induced by Cas9 was confirmed by sequencing and the Cas9 was segregated out (S1 Fig C).

Thus, we obtained three quintuple mutants: *dvl8 dvl11 rtfl9 rtfl11 dvl19, dvl1 dvl4 dvl8 rtfl11 dvl19*, and *dvl4 dvl8 dvl11 rtfl11 dvl19*. Again, the plants did not show noticeable defects in development when leaf morphology, inflorescence and floral morphology or fruit development were inspected (Fig 1A-E).

**Fig 1.**
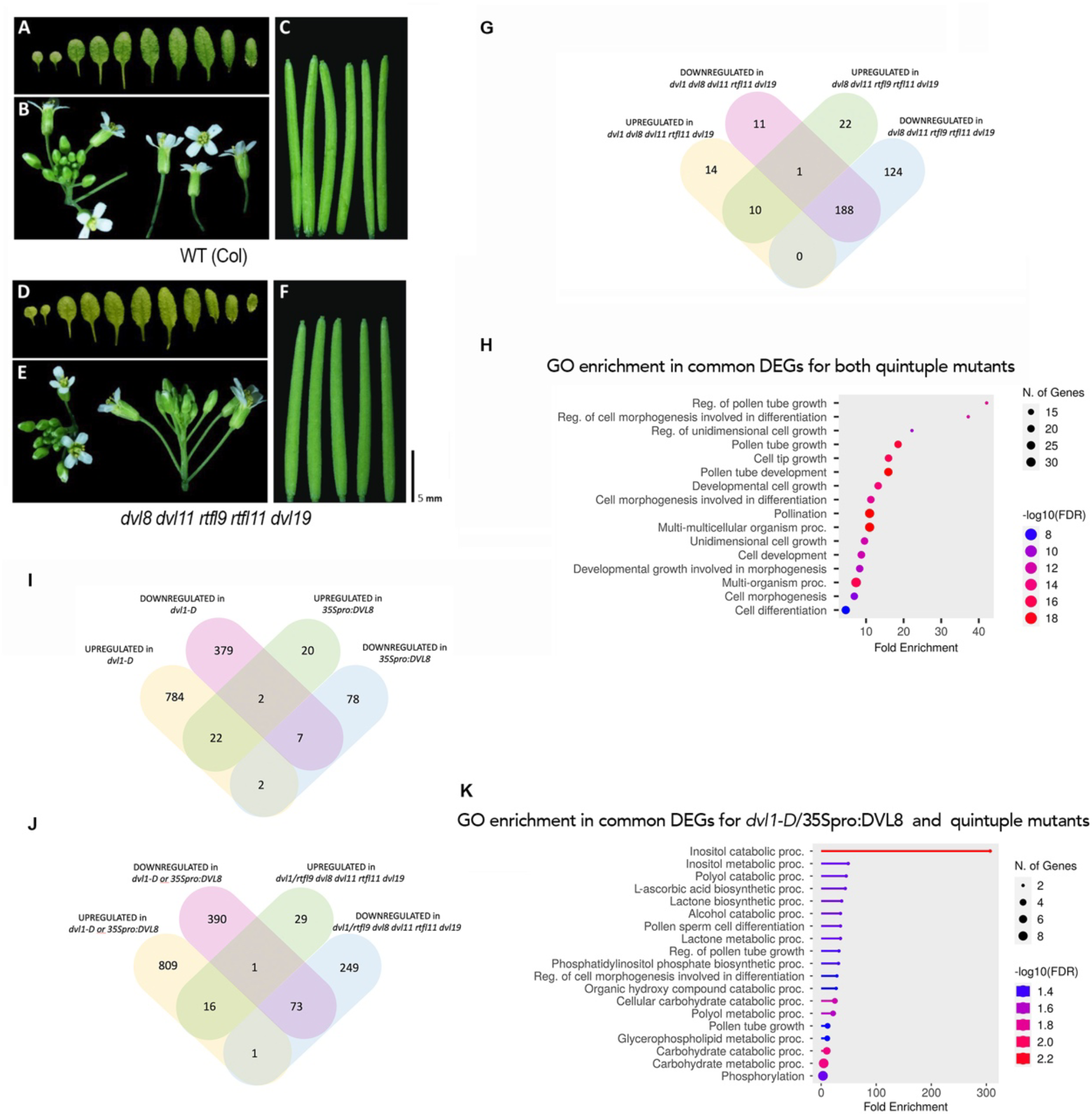
Phenotype of *dvl* quintuple mutants. (A-C) Morphology of wildtype plants. (D-F) Morphology of dvl quintuple mutants. (A, D) Rossette leaves of plants 15 days after germination. (B, E) Inflorescence of plants 10 days after bolting and of flowers at anthesis. (C, F) Fully elongated fruits from a single plant in equivalent positions. (G) Venn diagram shows DEGs shared between *dvl1 dvl8 dvl11 rtfl11 dvl19* and *dvl8 dvl11 rtfl9 rtfl11 dvl19* quintuple mutants. (H) GO analysis of the 199 DEGs common to both quintuple mutants. (I) Venn diagram shows shared DEGs between dvl1-D and 35S:DVL8 overexpression lines. (J) Venn diagram shows shared DEGs between overexpression lines and quintuple *dvl* mutants. (K) GO analysis of the DEGs common for at least one of the overexpression lines and one of the quintuple mutants.

The lack of visible phenotypes prompted us to compare the transcriptomic signatures of inflorescence apices of wildtype and mutant combinations (specifically *dvl1 dvl8 dvl11 rtfl11 dvl19* and *dvl8 dvl11 rtfl9 rtfl11 dvl19*). We collected whole inflorescences where the oldest flower was in anthesis in three independent biological replicates that were used for RNA sequencing. Transcripts with a log2 fold change (FC) >0.5 and <-0.5, and a p-adjusted value < 0.05 were considered as differentially expressed genes (DEG). Interestingly, and despite the absence of morphological defects and the unlikely role of DVL peptides in transcriptional regulation, significant changes in gene expression were caused by the mutation of DVL genes in both quintuple mutant backgrounds. We identified 224 DEGs in *dvl1 dvl8 dvl11 rtfl11 dvl19* vs. wildtype and 354 DEGs in *dvl8 dvl11 rtfl9 rtfl11 dvl19* vs. wildtype, where 199 were common to both experiments, most of them (188) downregulated in either mutant combination (Fig 1G).

With this list of DEGs we conducted a Gene Ontology (GO) analysis using the ShinyGO tool [14], focusing on the enriched terms in the category Biological Process. Only one, transmembrane transport, with four genes out of 10, was significatively enriched in the upregulated genes, while in the list of downregulated genes the most overrepresented categories were mainly related to processes like pollen tube development/growth and cell tip elongation /morphogenesis (Fig 1H). (S1 Table)

To complement these analyses, we included in the experiments the *dvl1-D* mutant, where *DVL1* is overexpressed by the insertion of a 4x35S enhancer element in the promoter of the gene, and a *35Spro:DVL8* line. Both lines share the typical phenotype associated to *DVL* overexpression, with small round leaves, protruding trichomes and short floral organs, although some differences are also noticeable, mainly related to fruit shape, that in *dvl1-D* plants have pointy valves in the apical-medial domain, while in *35Spro:DVL8* fruits are expanded laterally in the basal domain producing an arrow shape (S2 Fig). 1996 DEGs were identified when *dvl1-D* and wild-type inflorescences were compared, while only 131 DEGs appeared when *35Spro:DVL8* and wildtype inflorescences were compared, despite both lines showing similar phenotypic alterations in morphology. We also expected a substantial overlap on the DEGs in both experiments. However, only 33 DEGs were common to *DVL1* and *DVL8* overexpression, maybe reflecting the different effects of increasing *DVL1* expression by the insertion of the enhancer element in the endogenous *DVL1* promoter region or expressing *DVL8* more broadly with the 35S promoter (Fig 1I). These common DEGs were enriched in GO categories related to oxidative stress and responses, and also included three transcription factors, two of them members of the TCP family, TCP1 and TCP4, with well-characterized roles in the regulation of organ growth, and ZAT7, a member of the C2H2 family that negatively regulates growth in response to abiotic stress, and other genes involved in cell-wall remodeling functions (S2 Table) [15–19]. Interestingly, when inspected separately, the DEGs in 35Spro:DVL8 inflorescences respect to wildtype were enriched in several GO terms related to pollen development and pollen tube growth, whereas is the list of DEGs in *dvl1-D* inflorescences respect to wildtype, we found enriched mostly responses to different stimuli (S2 Table). These results suggested that, even likely working in a highly redundant manner, the different DVL peptides could participate in specific biological processes.

When comparing the effect of *DVL1* or *DVL8* overexpression with that observed in the quintuple mutants, only a reduced number of common DEGs were found (Fig 1J,). Surprisingly, most of these DEGs were similarly up- or down-regulated in all lines (both loss and gain of function, S3 fig), suggesting that an excess of DVL expression could have a dominant-negative effect and thus that at least part of the phenotypic defects found in the overexpression lines could be similar to those potentially caused by *DVL* loss of function. Again, these common DEGs were enriched in genes related to pollen development and pollen tube growth and, importantly, inositol metabolism, which is strongly related to cell elongation processes (Fig 1K) [20,21].

### DVL mutations reduce robustness of cell growth patterns

The overrepresentation of pollen-related categories among the DEGs in the transcriptomic analyses prompted us to check more carefully for developmental defects related to pollination in the mutants. At this point, a higher order combination of *dvl* mutations had been generated, so we used the septuple mutant *dvl1 dvl4 rtfl9 dvl8 dvl11 rtfl11 dvl19* for these analyses. The germination of pollen grains and the growth of pollen tubes *in vitro* was quantified and compared in wildtype and this septuple *dvl* mutant. The *dvl1-1D* line was also included. We did not observe significant differences in pollen germination among these genotypes after 2, 4, 6, or 8h (FIG 2A). Then, we measured the length of the pollen tubes 4 h after the grains were placed in the plates, since at this time point around 20% of the pollen grains or more had germinated in all samples and the length of the tubes was in a range that could be easily measured. While the average length of the pollen tubes was not significantly different among the three genotypes, we observed a greater dispersion of the data in the septuple mutant, where the pollen tubes had maximum-minimum lengths varying between aprox. 10 and 830 µm, in contrast with the wildtype that ranged between 10 and 425 µm (FIG 2B). *dvl1-D* pollen tube length had an intermediate dispersion, ranging from 15-650 µm. This suggested that DVL peptides could be providing robustness to pollen tube growth, perhaps in the context of a more general role in cell elongation.

To test whether this effect on uniformity of pollen tube growth influenced the fertilization of the ovules, we quantified the number of seeds produced by wildtype and *dvl* mutant plants. For this experiment, we used a higher order mutant, where additional DVL genes were mutated by CRISPR-Cas9: *dvl3 dvl4 dvl5 dvl6 rtfl9 dvl8 dvl11 rtfl11 dvl19,* which again did not show obvious alterations if development (S3 Fig). We collected fully developed siliques in positions 6 to 10 (from the base) in the main inflorescence of 12 individual plants of each genotype and scored the number of seeds per silique. Wild-type plants formed an average of 62 seeds per silique, within the range of 58-69, while the nonuple *dvl* mutants formed an average of 60 seeds, but within a wider range, 28-73 (FIG 2C-D). To test whether this effect could be affected by the position of the silique in the inflorescence or the overall state of individual plants, we quantified the number of seeds in siliques at positions 6-10 or 16-20 in 12 individual plants (Fig 2E). The increased dispersion of the data in the nonuple *dvl* mutant was similarly observed in all individuals, suggesting that is a general effect of *DVL* loss-of-function.

**Fig 2.**
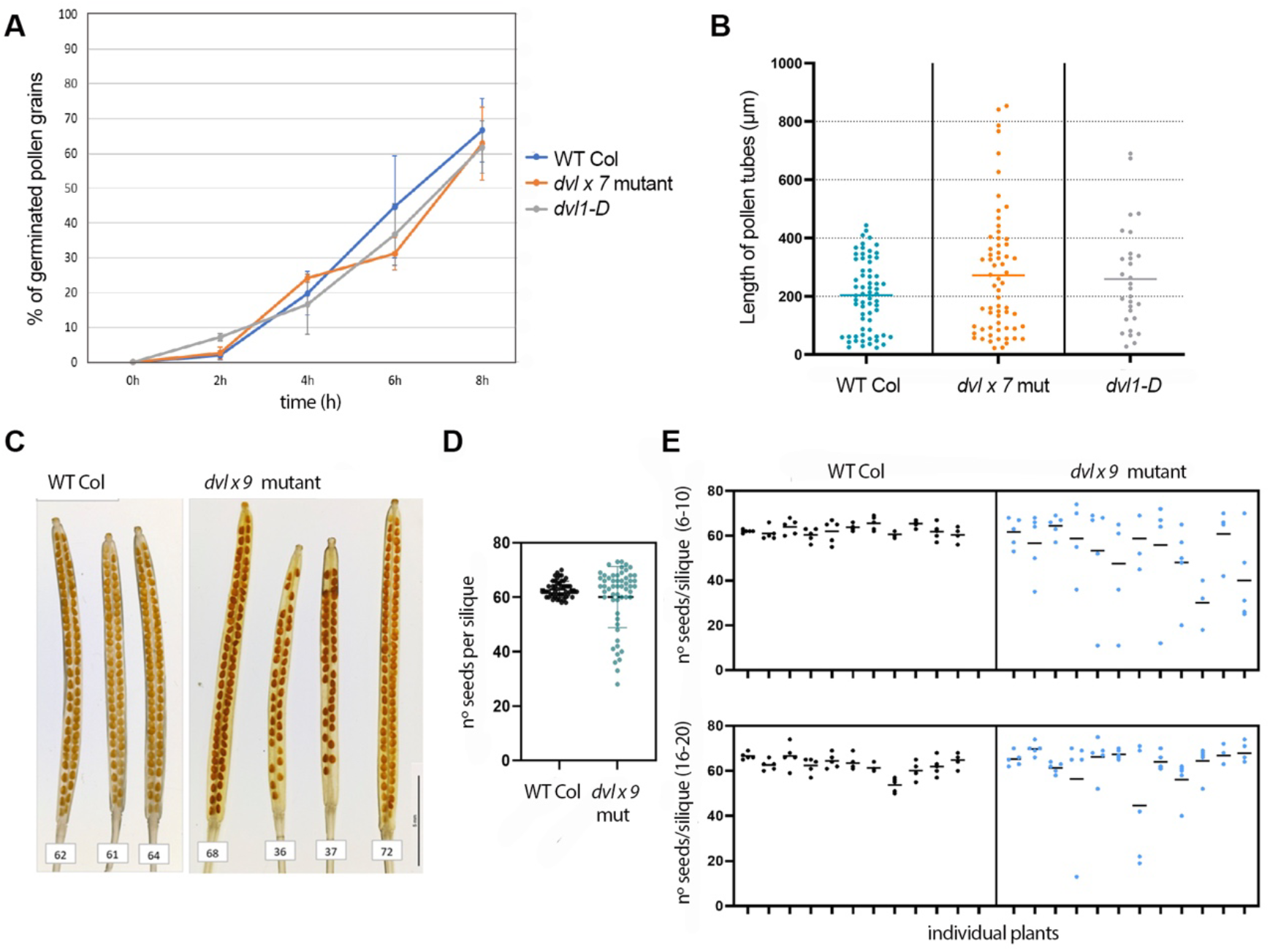
Effect of mutations in DVL genes on pollen tube elongation and seed production. (A) Percentaje of germinated pollen grains from wildtype, septuple *dvl1 dvl4 rtfl9 dvl8 dvl11 rtfl11 dvl19* mutants *(dvlx7),* and the dvl1-D overexpression line after being placed in plates containing germination media. No significant differences were observed. (B) Length of pollen tubes formed by pollen grains germinated in vitro 4 hours after being placed in germination media. Each individual dot corresponds to the length of a single pollen grain (C) Cleared siliques of wiltype (left) or nonuple *dvl* mutant (*dvlx9*, right) where the distribution of developed seeds can be observed. (D) Number of seeds per silique in wildtype and *dvlx9* plants. Individual dots correspond to the number of seeds in a single pod. Pods were pooled from diferent plants. (E) Number of seeds per silique collected from individual plants of wildtype and *dvlx9* backgrounds. Ten siliques per plant were collected in positions 6-10 (top panels) and 16-20 (bottom panels), where position 1 corresponds to the first silique formed in the main inflorescence.

To investigate whether the potential role of DVL peptides was restricted to pollen development or was more general in other cell elongation processes, we compared the growth of wild-type and septuple *dvl* mutant roots. The root apical meristem (RAM) plus the elongation zone (EZ) was determined by measuring the distance between the root tip and the first visible root hair and found to be significantly shorter in the mutants, suggesting that either cell elongation or cell division was affected (Fig 3A-B). While the number of roots hairs was not significantly different in the mutants (quantified as all root hairs from the most distal up to 2 mm above in the root; Fig 3D), root hair length (quantified for all root hais in these same 2 mm) was also reduced in the mutants and again showed a higher dispersion of the data, similarly to what we found in pollen tube growth (Fig 3 C, E). These results suggested that DVL peptides have a general role in providing robustness to cell elongation.

**Fig 3.**
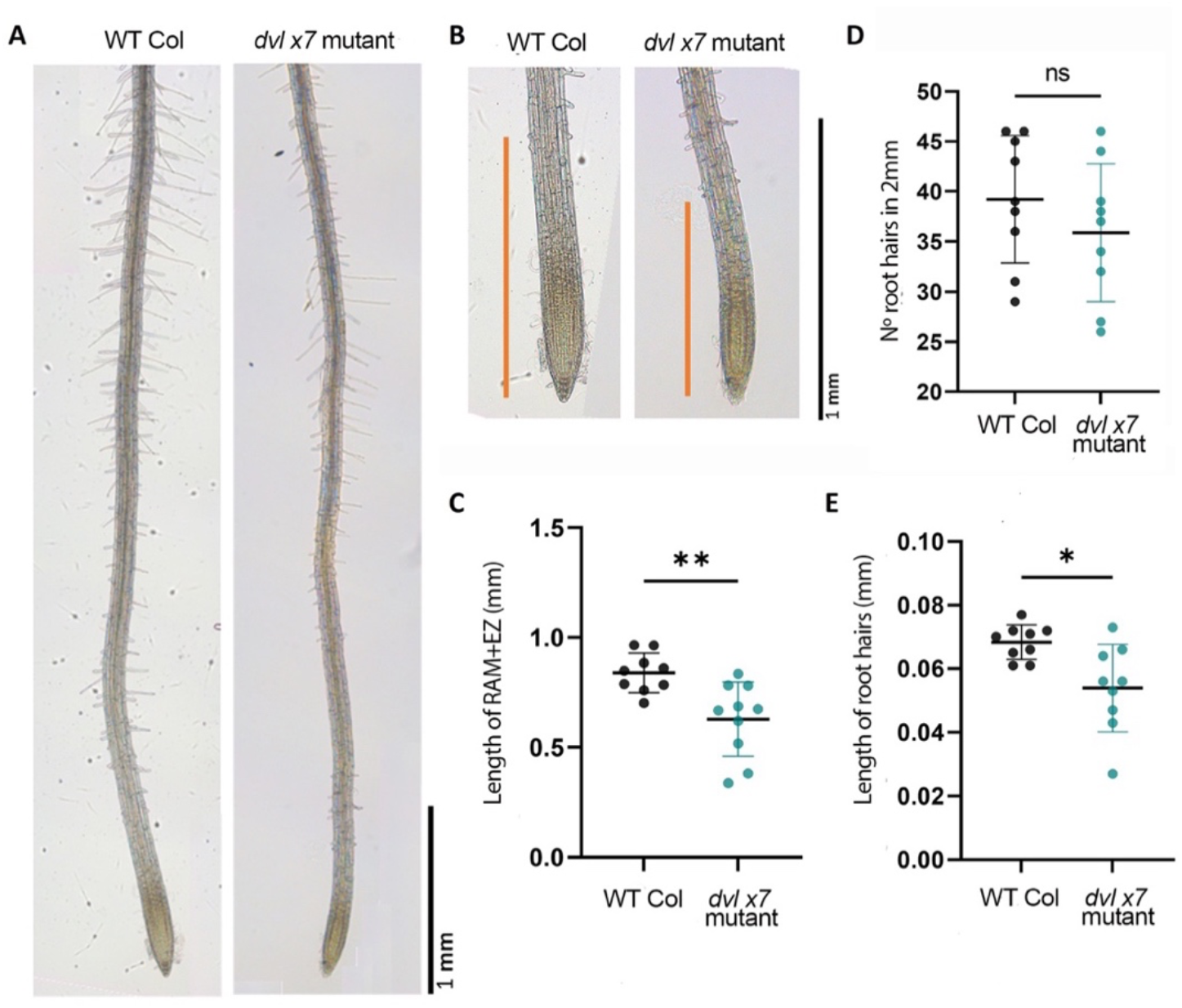
Effect of mutations in *DVL* genes in root development. (A) General morphology of roots from wildtype and *dvlx7* plants. (B) Close up of the root tips shown in (A). The orange bar spans the combined length of the root apical meristem (RAM) and the elongation zone (EZ), comprised between the most distal portion of the root meristem and the first observable root hair. (C) Length of the RAM+EZ in wildtype and *dvlx7* plants. (D) Number of root hairs in the most distal 2 mm of the differentiation zone of the root in wildtype and *dvlx7* plants. (E) Length of root hairs in the most distal 2 mm of the differentiation zone of the root in wildtype and *dvlx7* plants. Each dot represents the average value of all root hairs in an individual plant.

### DVL peptides might act as mobile elements

The subtle phenotypes associated to the combined loss of function of almost half of the DVL family genes suggested that DVL genes worked in a highly redundant manner. This could be the effect of active compensation mechanisms, where the mutation of one gene triggers the upregulation of other related genes. However, the most likely scenario was that of true genetic redundancy, since in the transcriptomic analyses of the quintuple *dvl* mutants we did not detect increased expression of the rest of the members of the family as a response to the loss of function of the mutated genes (S3 Table). To support this type of redundancy, the expression patterns of the different genes should overlap to some extent. In situ RNA hybridization experiments failed to detect DVL1, DVL8 or DVL11 expression in reproductive tissues, so we generated reporter lines where the predicted promoter of the corresponding genes drove the expression of the GUS reporter. The constructs also contained the 5’UTR region. We observed that DVL8pro:GUS, DVL11pro:GUS, RTFL9pro:GUS or DVL19pro:GUS constructs (all corresponding to genes of the same clade) conferred similar expression patterns to the reporter (Fig 4A, C; S4 Fig). In inflorescences GUS activity was detected in the vasculature of the stem, the pedicels, and floral organs. Expression was generally stronger in sepals and the filament of anthers, and also accumulated in the distal domain of the developing pistils, but was excluded from the stigmatic tissues once differentiated (Fig 4A, C; supp). DVL1pro:GUS lines also produced similar reporter activity, despite DVL1 belonging to a different clade (Fig 4E). In general, promoter activity was strong for all the genes tested, an unexpected result considering that in situ RNA hybridization experiments did not detect any signal, and the low expression of the different DVL genes in the inflorescence observed in the RNA-seq analyses. A possible explanation was that DVL mRNAs had a very rapid turnover. To test this hypothesis we also generated three additional lines: DVL8pro:DVL8-GUS, DVL11pro:DVL11-GUS and DVL1pro:DVL1-GUS, with the same promoter region used in the previous reporter lines and the 5’UTR, but including the coding region of the corresponding gene translationally fused to the GUS CDS. Surprisingly, the fusion protein was localized in different domains, almost complementary to those where the promoter was more active or, in the case of DVL1-GUS, almost undetectable in the inflorescence. DVL peptides do not have a predictable secretion signal and are unlikely to be processed, so these results suggest that either the mRNA contains elements in the coding sequence that promote long range movement or that DVL peptides are actively transported intercellularly.

**Fig 4.**
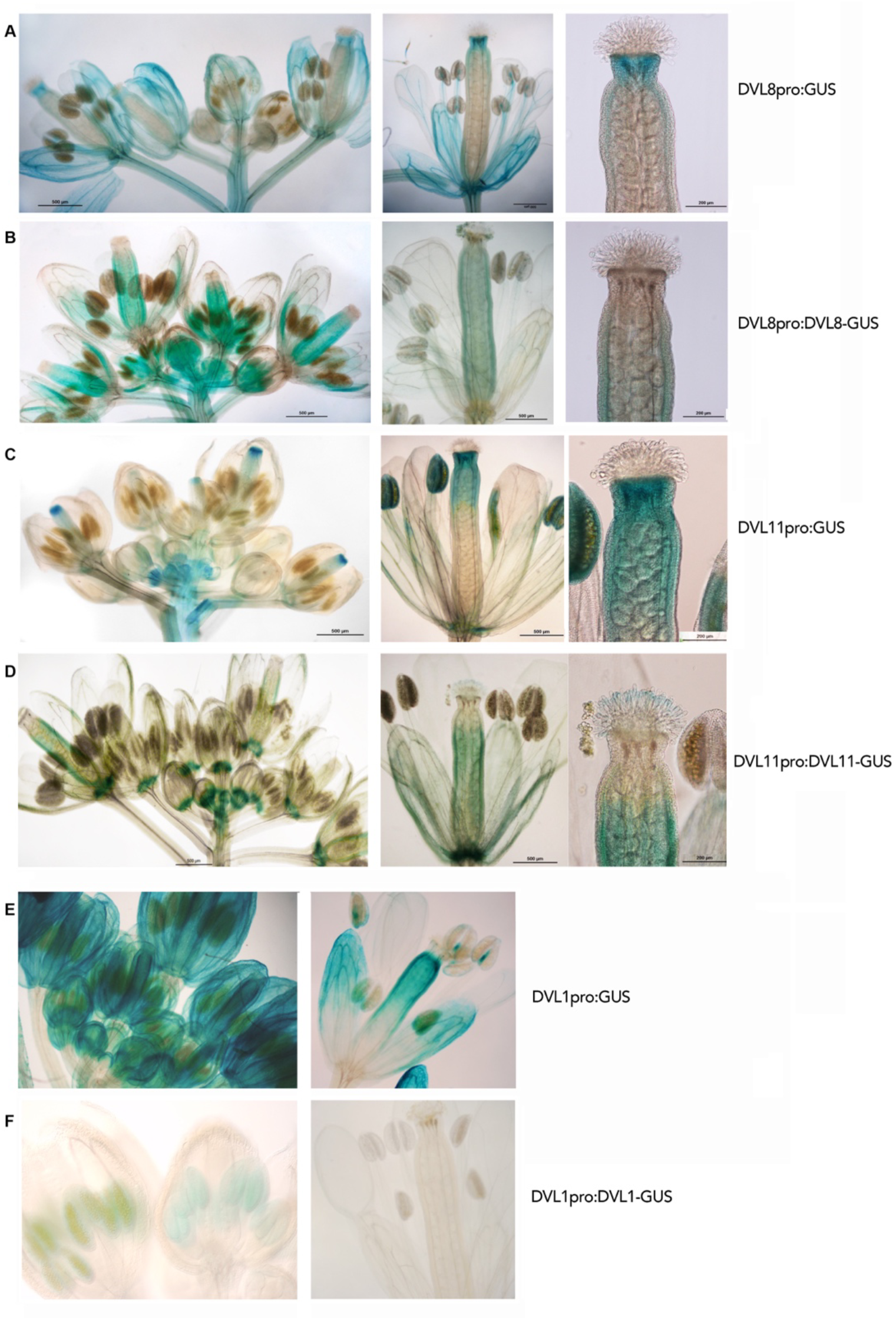
Activity of promoter:GUS or promoter:CDS:GUS reporters for different DVL genes. (A, C, E) GUS activity conferred by the promoter of DVL8 (A), DVL11 (C), and DVL1 (E) in reproductive tissues: inflorescences (left), florals organs as observed in flowers at anthesis (center) and in the apical zones of the gynoecium (right). (B, D, E) GUS activity conferred by the promoter of DVL8 (A), DVL11 (C), and DVL1 (E) and the promoter, 5’UTR and CDS of the corresponding genes, fused translationally to GUS in reproductive tissues: inflorescences (left), floral organs as observed in flowers at anthesis (center) and in the apical zones of the gynoecium (right).

## Discussion

The term “redundancy” refers to a situation in which there are more elements in a system than those needed to perform a function, and then the loss of one of these elements can be compensated by the others. In biological systems, a widely occurring form of redundancy is genetic redundancy, which frequently is derived from events of gene duplication, and plays a key role in evolution[22]. Functional redundancy provides robustness to processes and protects against deleterious mutations allowing beneficial ones to improve fitness or to evolve new functions, and then, true redundancy is usually limited to a small number of genes that have not undergone sub-functionalization or neo-functionalization yet [22–25]. The DVL family has gone through a rapid expansion in angiosperms, suggesting that some level of functional diversification must have occurred [9]. However, our study reports that even mutating almost half of the members of the family, ony subtle phenotypic defects are observed. One possible explanation is that the members of the DVL family in Arabidopsis represent an excepcional example of extensive genetic redundancy, and more genes need to be mutated to observe the consequences of the loss of DVL function. An alternative explanation is that DVL peptides play specific roles, at least for subclades within the family, but they remain hidden and only under specific conditions become relevant for developmental processes, stress responses, etc.

This reasoning has been the basis of our mutational strategy. We have first targeted a defined clade of five genes, based on the phylogenetic tree published by Narita et al.[6], assuming that the most closely related genes are also more likely to be redundant (Supp fig 2). The corresponding quintuple mutant *dvl8 dvl11 rtfl9 rtfl11 dvl19* did not present a conspicuous phenotype, somehow favoring the hypothesis of hidden roles in specific processes. To explore this idea, we chose to analyze the effect on the transcriptome of these combined mutations. This strategy had rendered good results in previous unrelated works [26], and in our case it has been of help to focus on particular processes that had been overlooked by just observing the overall morphology of the quintuple mutant. The differential expression analyses were useful to uncover the role of DVL genes in the elongation of pollen tubes and root hairs, where they might be conferring robustness to these processes. These phenotypes were subtle but clear and provide the basis for future studies on DVL mechanism of action. Our results also validate the use of transcriptomics to complement morphological observations in the characterization of genes of unknown function. It will be interesting to extend this strategy to higher order mutants and to analyze other tissues and/or treatments to uncover new aspects of DVL function.

In addition, by comparing the transcriptome of inflorescences of this quintuple mutant with that of a *dvl1 dvl8 dvl11 rtfl11 dvl19* combination, we could see differential effects on gene expression. Thus, the loss of function of all members of the clade had more impact on the transcriptome than maintaining the function of one member of the clade, *RTFL9*, and mutating *DVL1*, a gene belonging to a different group. This may suggest that the RTFL7-11 clade had redundant but specific functions as a subgroup, although we have to consider that this comes from a single experiment with samples restricted to inflorescences, and it is possible that it merely reflects statistic limitations of the analysis. More intriguing is the result of comparing the overexpression of *DVL1* and *DVL8*. The effects of the inflorescence transcriptome in both lines have few common features, despite the phenotypic similarities, supporting the idea of some degree of functional specificity of these two DVL peptides that belong to different clades. Again, an alternative explanation is that the enhancer element inserted the DVL1 promoter in *dvl1-1D* line and the 35S promoter driving the expression of DVL8 direct them to different expression domains and therefore cause differential effects in the transcriptome. However, independent studies have also analyzed the transcriptomic changes caused by overexpression of *DVL4* and *ROT4*, members of different subclades [6,27], and in the available lists of DEGs coming from these studies we also did not find extensive coincidences. Altogether, our work and these previous studies suggest that despite the likely high redundancy among the members of the family, there is some degree of functional specialization at least within different groups. Finally, when comparing the DEGs in overexpression and loss-of-function backgrounds, only a small proportion of DEGs were common to both situations and, importantly, these common DEGs were not regulated in opposite directions as it could be expected but responded in the same way to gain or loss of DVL function. The cases where overexpression of a wild-type gene mimics to some extent the phenotype of loss-of-function mutations are rare [28]. However, it is possible to find some examples, usually with proteins that form part of multiprotein complexes, and the excess of some components disrupt stoichiometry or interfere in some way with the correct functioning of the complex. For example, overexpression of yeast histone pairs *HTA1* and *HTB1* causes each the same transcription-related phenotypes as loss-of-function mutations in those genes [29,30]. To explore this possibility and to get more insights into the molecular mechanisms underlying DVL function it would be important to identify proteins that interact with DVL in the membrane or other subcellular locations.

Our work also has uncovered the unexpected movement of DVL, although it remains to be clarified whether it is mediated by RNA or protein transport and what is the functional relevance of this transport. In any case, it adds a new layer to the intriguing role of these molecules, that could be then regarded as signaling factors. In addition, if different DVL mRNAs/peptides are transported to different domains, at least in the floral tissues, it could explain why the overexpression with the same constitutive promoter (for example, CaMV 35S) causes the different fruit morphologies found for specific DVL genes [6,7,10,11].

Our study provides new data that may help to elucidate the molecular mechanisms mediating the proposed roles of DVL peptides in development and biotic/abiotic stress responses [6,10–12,31]. A common theme to all our experiments is the relationship of DVL peptides with pollen-related processes, but other works have also pointed to the responses to microorganisms [12,31]. However, we are still far from understanding these molecular mechanisms, the mode of action and the precise processes where DVL peptides may have a relevant function. Our work points to the participation of DVL peptides in processes related to cell elongation, where they may provide robustness, preventing uncoordinated growth or lack of uniform response to challenges eliciting cell wall modifications. At this point, it remains completely unknown how DVL peptides may participate in those processes and even if they are biologically relevant. It would be crucial to find interacting molecules that could shed some light on these putative functions and the mechanisms involved.

## Methods

### Plant material and growth conditions

The *Arabidopsis thaliana* accession Columbia (Col-0) was used as wild type. Overexpression lines *dvl-1D* and *35Spro:DVL8* reported in Valdivia et al. [10] were used in this report. Single T-DNA insertion lines (homozygous) were used for the different analysis and for generating multiple mutants by crossing: SALK_093633C (*dvl8*), SALK_204467 (*dvl11*), GK_321A11 (*rtfl11*) and SAIL_691_G09 (*dvl19*). CRISPR/Cas9 mutants and transgenic lines were generated as described below.

All Arabidopsis plants were grown on a 2:1:1 by volume mixture of sphagnum:perlite: vermiculite or Murashige and Skoog (MS) medium at 21°C under continuous light and long day (LD – 16 h light) conditions. The seeds for *in* vitro culture were surface-sterilized with 70% ethanol and 0.005% Triton-X-100 al 0.005% (v/v) for 3 min, then washed with 96% ethanol. To promote germination, seeds were stratified on soil at 4 °C for 3-5 days in the dark. Arabidopsis used for root growth studies were cultured and grown on MS medium in vertical direction.

### Plasmid construction and plant transformation

The reporter lines *DVL8_pro_:GUS*, *DVL8_pro_:DVL8:GUS*, *DVL11_pro_:GUS* and *DVL11_pro_:DVL11:GUS* were generated in this work. The *DVL8* promoter (2.8 kb), *DVL8* promoter fused to the *DVL8* CDS, *DVL11* promoter (3.6 kb) or *DVL11* promoter fused to the *DVL11* CDS were amplified and cloned in to the pCR8 vector using the pCR8/GW/TOPO TA Cloning Kit (Invitrogen). Then, these fragments were cloned into the destination vector pMDC163[32], which contains GUS by LR recombination (Invitrogen). In all cases, the integrity of all constructs was confirmed by sequencing.

The *dvl1*, *dvl4*, *rtfl9, dvl3*, *dvl5* and *dvl6* mutants were obtained following the published protocol by Wang et al. [13]. In the case of *DVL1*, *DVL4* and *RTFL9* one construction per gene was generated to make a big deletion of each gene, with two single guide RNAs (sgRNAs) sequence for each gene. In the case of *DVL3*, *DVL5* and *DVL6* only one sgRNAs sequence for each gene was introduced. T1 and T2 plants were screened for lesions at the corresponding DVL loci using Sanger sequencing and the sequences were analyzed using the *Synthego* web tool *ICE CRISPR Analysis* (https://www.synthego.com/products/bioinformatics/crispr-analysis).

Arabidopsis was transformed, in all cases, with Agrobacterium strain C58 pM090 using the simplified floral dip protocol [33]. Transgenic plants were selected on MS medium containing the selection method (20 mg/L hygromycin or 50 mg/L kanamycin). In CRISPR/Cas9 mutants, Cas9-free T3 seeds were selected by testing batches of seeds for hygromycin sensitivity.

### GUS staining

For **β** -Glucuronidase (GUS) histochemical detection, samples were treated for 15 min in 90% acetone and then washed with distilled water and incubated from to 16 h at 37 °C with GUS staining solution (50 mM sodium phosphate buffer, mM potassium ferrocyanide, 5 mM potassium ferricyanide, 1% Triton X-100 and 1 mM X-Glucuronic acid) in the dark. Then the stained samples were washed with water and clarified in series of ethanol solutions, and then observed under a Leica microscope (Leica DM5000) under bright-field conditions.

### Pollen germination and pollen tube growth assays

For pollen in vitro germination assay the protocol published by Smith and Wallace [34]was followed with minor modifications. Mature pollen grains were collected and spread on pollen germination medium containing 5mM CaCl_2_, 0.01% H_3_BO_4_, 5mM KCl, 1mM MgSO_4_,10mM, 10% sucrose (pH 7.5) and 1.5 % low melting agarose. After incubation in the darkness for 2, 4, 6 and 8h at 25°C, pollen germination was examined under a Leica microscope (Leica DM5000) under bright-field conditions. In vitro pollen germination frequency and in vitro pollen tube elongation rate was determined as previously described [34].

### Seed production analysis

For seed production in individual siliques, they were gathered individually after ripening but still green. Harvested siliques were placed in fixing solution (ethanol absolute, acetic acid (3:1) (v/v)) overnight at room temperature. The following day, fixing solution was exchanged by clearing solution (0.02 M chloral hydrate and 20% glycerol) and incubated overnight at room temperature. The cleared siliques were observed with a macroscope Leica DMS1000 under bright field conditions.

### Root tip and root hair observation

Lengths of root RAM and EZ regions and root hairs in 5-d-old seedlings grown on vertical MS medium without sucrose were measured under a Leica microscope (Leica DM5000). The data were analyzed using the ImageJ software (http://rsb.info. nih.gov/ij). To ensure the data from different samples are comparable, all the root hairs from the same root region (2.0 mm from the first root hair closed to root tip) were measured. Ten plants were randomly selected for measurement from each genotype.

### Quantitative RT-PCR analyses

Seedlings of 15 days-olds were harvested. RNA was extracted using the E.Z.N.A. Plant RNA Kit (Omega Bio-tek) and DNase treated with DNA free kit (Fermentas). RNA concentration and purity were verified using a NanoDrop Spectrophotometer ND1000 (Thermo Scientific). cDNAs were synthesized from 2 to 3 μg of total RNA using SuperScript IV and Oligo dT12-18 (Thermofisher) following manufacturer’s instructions. The RT-qPCR was performed in QuantStudio™ real time PCR system (Thermofisher) and used *Premix PyroTaq Eva Green qPCR Mix Plus* (GMC) to monitor double-stranded DNA synthesis. The Ct value was obtained from an automatic threshold. Results were normalized to the expression of the *ACT* reference gene. The 2^−ΔCC(T)^ was shown as relative expression level. Three biological replicates were performed for each sample.

### RNA isolation and RNA-Sequencing analysis

RNA for RNA-seq was extracted with the RNeasy Plant Mini Kit (Qiagen) from inflorescences and flowers in post-anthesis, adding the DNase to the column with the RNase-Free DNase Set (Qiagen) according to manufacturer’s instructions. RNA concentration and purity were verified using a NanoDrop Spectrophotometer ND1000 and a Bioanalyzer 2100 (Agilent Technologies). The cDNA libraries preparation, sequencing and analysis were made by Novogene (UK) Company Ltd. using Illumina Novaseq6000 system with a PE150 strategy (pair end – 150 pb simple ends).

For the bioinformatic and statistical analysis, the input files were processed using the *cutadapt* software (http://cutadapt.readthedocs.io/en/stable/guide.html). RNA-seq reads were aligned to the reference genome of Arabidopsis available at the TAIR database (http://arabidopsis.org; Lamesch et al., 2012) using *hisat2* software (https://ccb.jhu.edu/software/hisat2/index.shtml). The abundance estimation of the transcripts was performed using the *htseq-count software* (https://htseq.readthedocs.io/en/release_0.11.1/count.html), and the differentially expressed transcripts were estimated using *DESeq2* package (https://bioconductor.org/packages/release/bioc/vignettes/DESeq2/inst/doc/DESeq2.html). The gene abundance is represented by RPKM value (Reads Per Kilobase of exon model per Million mapped fragments). To identify the differentially expressed genes Fold Change was calculated dividing the problem sample RPKMs between control sample RPKMs and this Fold Change was transformed to a log2., and statistical tests were applied. The selection of differentially expressed genes were made with the following criteria: RPKM >0.5, *log. 2-fold change* >0.5 o <-0.5 y un *p*-value adjusted < 0.05.

The enrichment analysis of gene ontology categories (GO terms) was performed using the ShinyGO database and web tool (http://bioinformatics.sdstate.edu/go/; Ge et al., 2020).

## Acknowledgements

The authors would like to thank Javier Forment (Bioinformatics, IBMCP) for help with transcriptomic analyses. We also thank Francisco Madueño (IBMCP) for critical reading of the manuscript and members of Ferrandiz lab for discussions and suggestions.

## Supporting material

**S1 fig. Generation and characterization of mutants for DVL genes.** (A) T-DNA insertion lines used in this study were obtained from NASC or, in the case of RTFL9, obtained by Crispr-Cas9 genome editing. These mutants were combined and used in the transcriptomic analysis described in the main text (B) Expression of the corresponding genes in the T-DNA insertion lines shown in panel A. (C) Additional mutants generated by Crispr-Cas9 genome editing. Orange triangles indicate the confrimed position of the T-DNA insertion; green bars indicate the position of the RNA guides used for editing; orange bars indicate the deleted portion of the gene in the edited lines used for this study. For each gene, the domains corresponding to the 5’ – and 3’-UTR regions are indicated, as well as the coding region (light blue) and the conserved DVL/RTFL domain (purple).

**S2 fig. Phenotype of the DVL overexpression lines used in this study.** Inflorescence and mature fruit morphology are shown for *dvl1-1D* (top panels) and 35S:DVL8 lines (bottom panels)

**S3 fig. Phenotype of *dvl* nonuple mutant.** (A-C) Morphology of wildtype plants. (D-F) Morphology of dvl nonuple mutants. (A, D) Rossette leaves of plants 15 days after germination. (B, E) Inflorescence of plants 10 days after bolting and of flowers at anthesis. (C, F) Fully elongated fruits from a single plant in equivalent positions.

**S4 fig. Expression pattern conferred by DVL19 and RTFL9 promoters.** GUS activity in flowers around anthesis stage conferred by the promoters of DVL19 (left) or RTFL9 (right).

**S1 Table. GO term enrichment in DEGs in dvl mutant inflorescences.**

**S2 Table. GO term enrichment in common DEGs for *dvl* mutants and DVL overexpressing inflorescences.**

**S3 Table. Expression of *DVL* genes in inflorescences of the quintuple *dvl* mutant combinations.** Data was retrieved from the results of the RNAseq experiments described in the main text. Table shows RPKM (Reads Per Kilobase Million) corresponding to each DVL/RTF gene in the inflorescences WT, *dvl1 dvl8 dvl11 rtfl11 dvl19* and *rtfl9 dvl8 dvl11 rtfl11 dvl19* mutants. Red asterisk indicates that the gene is mutated in the corresponding sample.

